# Genetic mechanisms and biological processes underlying host response to ophidiomycosis (Snake Fungal Disease) inferred from tissue-specific transcriptome analyses

**DOI:** 10.1101/2022.03.25.485740

**Authors:** Samarth Mathur, Ellen Haynes, Matthew C. Allender, H. Lisle Gibbs

## Abstract

There is growing concern about infectious diseases in wildlife species caused by pathogenic fungi. Detailed knowledge exists about host pathology and the molecular mechanisms underlying host physiological response to some fungal diseases affecting amphibians and bats but is lacking for others with potentially significant impacts on large groups of animals. One such disease is ophidiomycosis (Snake Fungal Disease; SFD) which is caused by the fungus *Ophidiomyces ophidiicola* and impacts diverse species of snakes. Despite this potential, the biological mechanisms and molecular changes occurring during infection are unknown for any snake species. To gain this information, we performed a controlled experimental infection of captive Prairie rattlesnakes (*Crotalus viridis*) with *O. ophidiicola* at different temperatures. We then generated liver, kidney, and skin transcriptomes from control and infected snakes to assess tissue specific genetic responses to infection. Given previous SFD histopathological studies and the fact that snakes are ectotherms, we expected highest fungal activity on skin and a significant impact of temperature on host response. In contrast, we found that most of the differential gene expression was restricted to internal tissues and fungal-infected snakes showed transcriptome profiles indicative of long-term inflammation of specific tissues. Infected snakes at the lower temperature had the most pronounced overall host functional response whereas, infected snakes at the higher temperature had overall expression profiles similar to control snakes possibly indicating recovery from the disease. Overall, our results suggest SFD is a systemic disease with a chronic host response, unlike acute response shown by amphibians to *Batrachochytrium dendrobatidis* infections. Our analysis also generated a list of candidate protein coding genes that potentially mediate SFD response in snakes, providing tools for future comparative and evolutionary studies into variable species susceptibility to ophidiomycosis.

**Author summary:** Ophidiomycosis (Snake Fungal Disease; SFD) is an infectious fungal disease in snakes that has been documented in more than 40 species over the past 20 years. Though many snake species seem vulnerable to SFD, little is known about how snake physiology changes in response to infection with the causative fungus, *Ophidiomyces ophidiicola*. Here we report results from the first experimental transcriptomic study of SFD in a snake host. Our goals were to identify genes with a putative role in host response, use this information to understand what biological changes occur in different tissues in snakes when infected with *O. ophidiicola*, and determine if temperature has an impact in these ectothermic animals. We conclude that SFD is a systemic disease with a chronic inflammation leading to deterioration of internal organs and that these physiological impacts are more pronounced at low rather than high temperatures. These results contrast with fungal infections in amphibians where hosts show an acute response mostly restricted to skin. Our list of candidate genes carry utility in potentially diagnosing genetic susceptibility to SFD in snake species of conservation concern.

## Introduction

Infectious wildlife diseases are an increasing threat to global wildlife diversity and have led to significant species declines as well as impacts on human health and livestock (1, 2). One class of wildlife infectious diseases are caused by pathogenic fungi. Two well-studied examples of fungal diseases in wildlife are chytridiomycosis in amphibians caused by the chytrid fungus *Batrachochytrium dendrobatidis* (*Bd*) (3), and white-nose syndrome in bats caused by fungus *Pseudogymnoascus destructans* (4). Infection with *Bd* disrupts the integrity of skin and is responsible for amphibian species declines worldwide (5, 6). Skin is a critical organ for amphibians and is involved in physiological activities such as respiration, ion balance, hydration, and defense against other pathogens; thus, infected hosts have high mortality (7, 8). White-nose syndrome is one of the most damaging infectious disease epidemics in bats (9) that caused extirpations of entire populations of many bat species (9, 10). The lesions caused by *P. destructans* are mostly found on bat ears and nose, but infections are most severe in the wing and tail tissues. *P. destructans* infection disrupts crucial regulatory functions like thermoregulation, gas exchange and immune function (11) which subsequently results in loss of fat store and starvation and eventual death of the bat host (12).

Numerous studies on both diseases (9, 13) have revealed detailed information about the effects of infection on different tissues (14), the effects of key environmental factors like temperature (15) and humidity (16), the role of co-infections with other pathogens (14), and even variable immune responses (17) and susceptibility among different species (18). This information is crucial to understand the disease systems, prevent outbreaks, and identify vulnerable populations. Fungal pathogens have been recognized in numerous additional taxa, including sea turtles (19), corals (20), lizards (21), honeybees (22), and various plants (23, 24), but estimates of disease prevalence and understanding of host responses to such pathogens are lacking in most species (25–27). The recent emergence and global spread of fungal pathogens have led to increased surveillance efforts for known pathogens and the need to understand how specific diseases impact infected individuals to understand the pathological mechanisms that underlie disease infections (23, 25, 27, 28)

One example of a poorly understood but potentially impactful disease is ophidiomycosis (snake fungal disease; SFD). SFD is a recently identified fungal disease in snakes caused by the fungus *Ophidiomyces ophidiicola* and has been detected in many free-ranging and captive snake species (29–32). *O. ophidiicola* is a generalist fungus and is known to infect a wide range of snake species with different ecologies irrespective of taxonomy and habitat (33). First reported in the mid-2000s, but likely present as early as the 1940’s (34–36), SFD has since been documented in more than 30 species of wild snakes in the United States and Europe (31) and instances of SFD have also been reported in Australia (37) and South East Asia (38, 39). Clinical signs of ophidiomycosis vary among individuals, from general signs such as lethargy, skin lesions, excessive shedding, to crusts, granulomas, corneal opacity, and ulcers on the head and body in more severe cases (29). Ophidiomycosis has the potential to cause widespread morbidity and mortality in snakes (31, 32, 35, 40, 41) but the mode of infection, specific mechanisms of pathology, and physiological responses by infected individuals are unclear despite the value of this information for understanding and mitigating the impact of this disease (29, 42).

Specifically, the two key elements of SFD pathology that are still unknown are (a) Is the snake host response is localized to the skin, as in the case of Bd infections in amphibians, or whether SFD is more systemic disease, like the white nose syndrome in bats? and (b) What is the influence of temperature on disease severity and host response of infected individuals? Experimental transcriptomics offers and approach to address these questions by allowing us to identify differentially expressed genes in multiple tissues, and then using functional enrichment and pathway networks approaches to identify what biological processes are occurring differently among tissue of infected and uninfected hosts.

Previous histopathological studies (43) identified *O. ophidiicola* infection to be localized to skin (30), so we expect the highest fungal activity on skin. Therefore, if the snake response to fungal infection is similar to amphibians, we predict most differential expression on skin tissue (14). Secondly, since snakes are ectotherms i.e., their thermoregulation is dependent on external temperatures, we predict that temperature would have a crucial impact on overall host response. Field evidence suggest that many infected snakes preferably move towards higher temperatures (44, 45) and can potentially recover from SFD (46). Additionally, pathogenic activity of many fungal species is temperature dependent and host defense against the infection is more effective at higher temperature in many vertebrate species (47, 48). Studies of SFD in free ranging snakes have also indicated that SFD severity declines with higher temperatures and higher fungal prevalence in cooler seasons (45).

Here, we studied the snake host response to SFD by performing controlled *O. ophidiicola* exposure experiments in Prairie rattlesnakes (*Crotalus viridis*) at 20⁰C and 26⁰C. Prairie rattlesnakes are a common and widely distributed snake species and are closely related to many species that are susceptible to SFD (33) which makes them a good model for controlled exposure experiments. We describe the genetic and physiological changes in multiple tissues due to *O. ophidiicola* exposure and also the effect of different temperatures by comparing the changes in transcriptome profiles of different organs under different conditions. Finally, we identified a list of candidate genes that are putatively involved in host response which could be used in diagnostic screening of more vulnerable populations and species of snakes, as done in *Bd* and other wildlife diseases (49).

## Results

### Experimental infections, transcriptome sequencing, and data pre-processing

All snakes were free of clinical signs over 24 months prior and at the start of the study. First clinical signs in *O. ophidiicola* infected snakes were observed around 40-45 days post infection (dpi). All infected snakes at 20°C were euthanized before the end of the study due to the severity of clinical signs, while only one infected snake was prematurely euthanized at 26°C. The remaining two infected snakes at 26°C survived with mild-to-moderate clinical signs untill the end of the study (90 dpi). All uninfected control animals survived throughout the study period (Table S1).

RNA was isolated and sequenced from liver, kidney, and skin tissues collected from each infected and uninfected snakes at the end of the study. Following sample processing and evaluation of library sequence quality, we retained transcriptome sequence data from 10 liver, 11 kidney, and 10 skin tissues from snakes subjected to different temperature (26°C vs. 20°C) and infection status (infected vs. control) (Fig. 1A; Table S1). Our sequencing of the RNAseq libraries resulted in the generation of a mean of 31.7 million read pairs (SD = 16.8 million) per sample (Table S2). After adapter trimming and removal of low-quality reads, the overall alignment rate for the remaining filtered reads to the *C. viridis* reference genome for each sample was 59.3% ± 14.8% (mean ± SD; Table S2) with 8.3% ± 2.3% of all aligned read pairs successfully assigned to the annotated regions of the *C. viridis* assembly. We measured alignment rate as the percentage of transcripts that mapped uniquely and concordantly to the reference genome. The alignment percentage to the *C. viridis* reference genome was lowest in skin tissues (liver = 61.6 ± 6.2%; kidney = 65.0 ± 3.7%; skin = 50.8 ± 23.3%) and one sample had only 1% of total transcripts aligning (Table S2).

**Fig 1.**
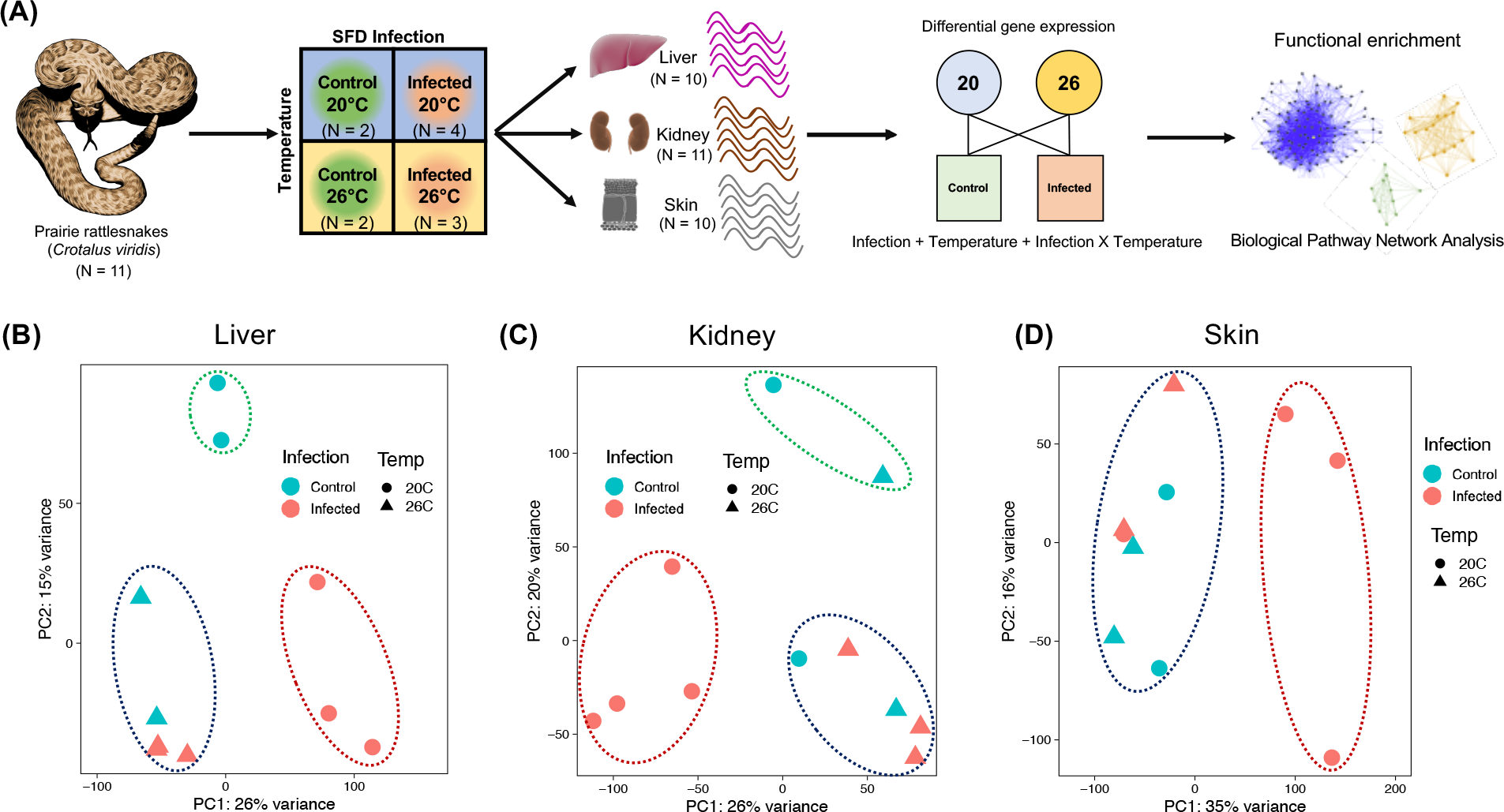
Methodological overview of the study and overall individual gene expression profile. (A) Schematic showing the experimental design for the controlled ophidiomycosis infection trial to study the host response. Prairie rattlesnakes (*Crotalus viridis*) were exposed to the causative fungus *Ophidiomyces ophiodiicola* (“infected”) or sham inoculations (“control”) at either higher (26°C) or lower (20°C) temperatures. N = number of samples within each group. After the end of the study (90 dpi), RNA was extracted and sequenced from liver (N=10), kidney (N=11), and skin (N=10) tissues of each individual. Differential gene expression (DGE) analysis was performed using infection status, temperature, and the interaction between infection and temperature (Infection X Temperature) as fixed effects. Each tissue (liver, kidney, and skin) was analyzed independently. Functional enrichment and pathway networks were then inferred from biological functions of differentially expressed genes (DEGs). (B-D) Tissue specific expression profile and individual clustering based on normalized read count data of all expressed genes in Liver, (C) Kidney, and (D) Skin. *O. ophidiicola* infected snakes at the lower temperature (red circles) cluster independently and are separated on PC1 axis (x-axis) in all three cases. Infected snakes at the higher temperature (red triangles) have similar profile as control snakes at the higher (blue triangles) or the lower (blue circles) temperature. The dotted circles represent the number of clusters identified using a discriminant analysis.

### Overall host response to SFD exposure is temperature dependent

We first visualized the sample groupings based on overall gene expression data using multivariate analyses. PCA plots generated from log normalized read counts of all assembled transcripts indicate that infected snakes at the lower temperature had an expression profile that differed from all other tissue-treatment combinations in all three tissue types (Fig. 1B-D). Both hierarchical clustering and discriminant analysis on principle components (DAPC) clustered infected snakes at 20⁰C as one group with all other samples forming separate groups (Fig. 1B-D, Fig. S1). More interestingly, infected snakes maintained at 26⁰C show a similar overall gene expression profile as all the control snakes. This means that by the end of the study, host response to *O. ophidiicola* infection is different from controls at the lower but not at the higher temperature treatment. Even though infected samples clustered based on temperature, no such pattern existed for control samples as different control samples clustered together, regardless of temperature conditions (Fig. 1B-C; Fig. S1). Overall, these results indicate that snakes infected with *O. ophidiicola* at low temperature respond most differently with a unique expression signature, and that high similarity in gene expression between infected snakes at the higher temperature and control snakes by the end of the study is indictive of possible recovery.

### Fungal gene expression identified only in skin tissues

To identify if fungal transcripts were present in our RNASeq data, we mapped the aligned filtered reads to the *O. ophidiicola* reference genome (see Methods). We expect that tissues with more actively growing fungus would have greater expression of fungal transcripts and thus, would have a higher alignment rate to the *O. ophidiicola* reference genome. We observed that most of the fungal transcripts in our samples were identified in skin tissues at the lower temperature (29.8% ± 37.4%) whereas all other samples (including skin tissues at the higher temperature treatment) had < 0.5% fungal transcripts (Fig. S2). We next identified expressed fungal genes based on highest transcript counts (i.e. highest depth of coverage). In each skin tissue, we observed peaks of high expression at the same region of the fungal genome (see e.g. in Fig. S3). Since the *O. ophidiicola* reference genome is not yet annotated, we extracted the reference sequence corresponding to the highest transcript peaks and used BLAST (https://blast.ncbi.nlm.nih.gov/) to identify homologous genes in other fungal genomes. We identified expression of genes than encode myosin class V proteins (identity 80.1%, e-value = 0), chaperone protein DnaK (identity 86.6%, e-value = 0), and beta-glucosidase 4 (identity 77.1%, e-value = 2e-15). Myosin proteins are highly conserved in fungi (50) and all chytrid species contain at least one myosin class V protein (51) that have a function in intracellular transport.

They are shown to localize in the actively growing hyphae (52). Similarly, beta-glucosidase enzymes breakdown complex macromolecules like cellulose or keratin (53) so, they might be important for damage to host skin tissues, and DnaK is part of the Heat Shock Protein (HSP) protein complex shown to be active during pathogenesis (54). Taken together, these results indicate that only skin tissues at the lower temperature have fungal expression and the fungal genes that have highest expression are likely involved in fungal growth and infection.

### SFD exposure leads to greater differential gene expression in internal organs compared to skin

We conducted differential gene expression (DGE) analyses from the counts of all assembled transcripts for each tissue separately (Fig. 2A-C). Our analysis design included temperature (“Temperature”; low = 20°C vs. high = 26°C), infection status (“Infected”; *O. ophidiicola* infection vs. control), and an interaction effect between temperature and infection (“Infected X Temperature”) as fixed effects (Fig. 1A). We identified a total of 776 DEGs in liver (Infected = 503; Temperature = 182; Infected X Temperature = 91; Fig. 2A), 705 DEGs in kidney (Infected = 507; Temperature = 131; Infected X Temperature = 67; Fig. 2B), and only 17 DEGs in skin (Infected = 15; Temperature = 2; Fig. 2C). We did not identify any DEGs due to the interaction between infection and temperature in skin. In terms of changes in expression, the majority of genes in liver (457/776; 58.9%) and skin (12/17; 70.6%) were upregulated, whereas majority of the genes in kidney (452/705; 64.1%) were down regulated.

**Fig 2.**
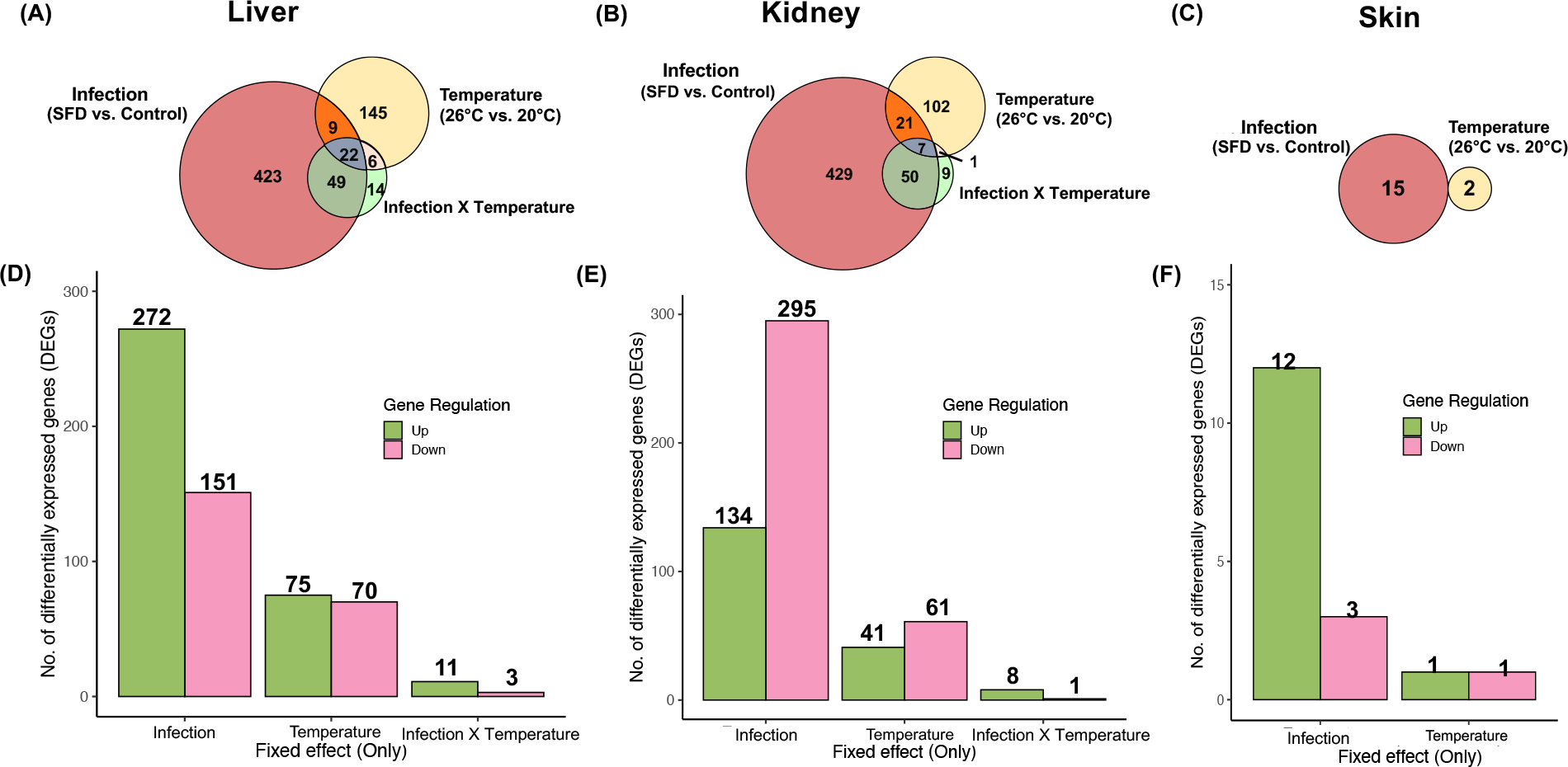
Differential gene expression (DGE) due to fixed effects. Euler plots showing overlap in differential gene expression due to fixed effects of *O. ophidiicola* infection (Infected vs. Control), temperature (26°C vs 20°C), and the interaction between infection and temperature (Infection X Temperature) for (A) liver, (B) kidney, and (C) skin tissues. The number within each plot represent the number of differentially expressed genes. There was no interaction between infection and temperature in the analysis of skin tissue. Panels D-F show the number of upregulated (green) and downregulated (pink) genes unique to each fixed effect. There were more upregulated than downregulated genes for each effect in (D) liver, whereas more genes were downregulated due to infection and temperature in (E) kidney. More genes were upregulated in (F) skin due to infection.

To identify DEGs due to the impact of a single fixed effect, we isolated DEGs that did not overlap with the two other fixed effects. These results are shown in Fig. 2D-F and Fig. S4. In all three tissue types, *O. ophidiicola* infection alone caused more changes in gene expression than either temperature or the interaction between temperature and infection (liver = 423; kidney = 429; skin = 15; Fig. 2D-F). Most of the DEGs due to infection only were specific to each tissue (Fig. S4) which indicates that *O. ophidiicola* infection impacts different biological processes in different internal organs, specifically in liver and kidney.

### SFD exposure induces a pro-inflammatory response and disrupts metabolism in liver

Most of the upregulated genes in liver were enriched in Biological Processes Gene Ontology (GO:BP) terms, “cellular response to organic stimulus” (35%; 475/1369), “developmental/metabolic processes” (49%; 666/1369), “immune response” (6%; 76/1369) and “cell death/apoptosis” (9%; 116/1369) suggesting positive cell differentiation as an inflammatory immune response to organic antigens (Fig. 3A). Upregulated genes like Colony-stimulating factor 1 receptor (*CTF1R*) mediate macrophage signaling, and *CTF1R* expression is crucial for the differentiation and survival of the mononuclear phagocyte system (55). In contrast, *MKNK2* is associated with interleukin-1 signaling pathway and its overexpression is associated with cell proliferation and reduction of apoptosis (56). We also identified 11 upregulated genes that are involved in the MAPK signaling pathway that initiates from a diverse range of stimuli and elicit an appropriate physiological response including cell proliferation, differentiation and migration, development, inflammatory responses and regulation of apoptosis (Fig. S5). Our results indicate that *O. ophidiicola* infection triggers an immune response in the liver and facilitates an inflammatory response by positively regulating cell proliferation and inhibiting programmed cell death.

**Fig 3.**
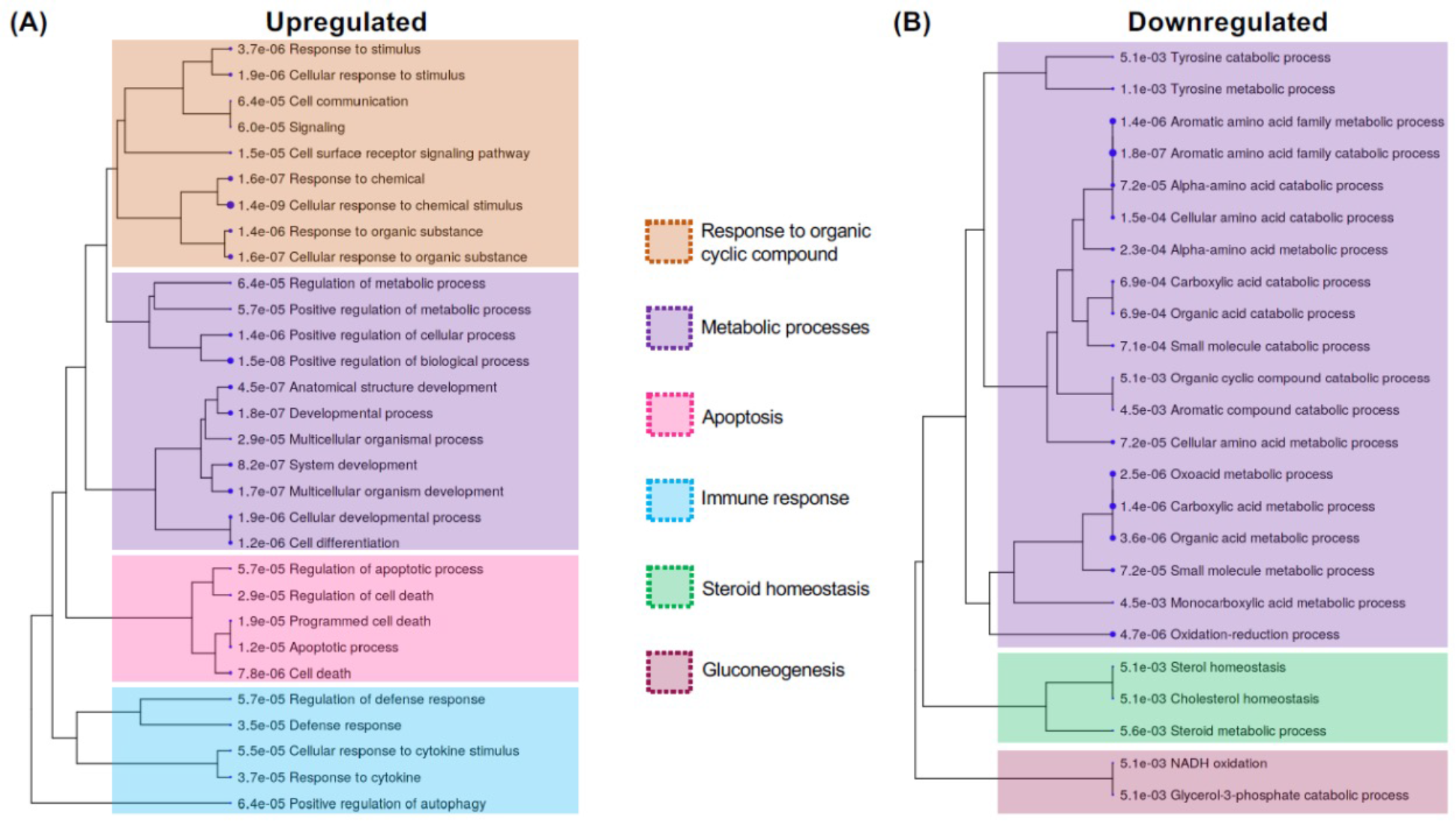
Biological processes that were differentially regulated in liver due to SFD exposure. Hierarchical clustering tree summarizing the correlation among significant Biological process gene ontology terms (GO:BP) for genes that were (A) upregulated, or (B) downregulated in liver. Numbers indicate p-values after FDR correction for multiple testing. Each process terms were categorized as subclass of specific parent terms as represented by colored boxes.

In terms of DEGs that were downregulated in liver tissue, most of the genes represent amino acid metabolism (91%: 182/200), steroid homeostasis (7%; 14/200), and gluconeogenesis (2%: 4/200; Fig. 3B). Steroid hormones mediate stress induced metabolic regulation and immune modulation (57). Negative regulation of steroid homeostasis in liver is associated with a pro-inflammatory response (57) and reduction of protein metabolism for glucose and glycogen synthesis (58). This means that the inflammatory response to *O. ophidiicola* results in liver being unable to metabolize proteins necessary for proper functioning. We also created networks or clusters of co-expressed genes (“modules”) using weighted gene co-expression network analysis to test if any modules were significantly associated with infection. After adjusting P values to account for multiple testing, we identified five modules significantly associated with infection status in liver tissue (Table S3) including genes enriched in Peroxisome KEGG pathway (Fig. S6). Peroxisomes break down complex fatty acids and subsequently regulate multiple metabolic pathways (59). Most of the genes within the peroxisome pathway were downregulated in liver tissues from infected snakes (Fig. S7). Peroxisome activation genes like *PPARα* downregulates various immunity-related pathways (59), so the lower production of *PPARα* in liver tissue from infected snakes is a signature of higher immune response.

Overall, our DGE analysis indicates that *O. ophidiicola* infection is associated with stress induced liver inflammation, as well as lower steroid production and protein metabolism which leads to lipid and protein accumulation within hepatic cells. These physiological changes are hallmarks of chronic liver diseases like liver injury and fatty liver disease and are characterized by fibrosis and cirrhosis of the liver (60).

### SFD exposure lowers protein metabolism and disrupts ion balance in kidney

In kidney tissues from infected snakes, most of the overexpressed genes reduce protein metabolism by inhibiting proteolysis, and protein degradation, and catalytic activity in kidney cells (Fig. 4A). Proteolysis eliminates irregular proteins, controls cellular regulatory processes, and provides amino acids for cellular remodeling (61). This means that in kidney tissues from infected snakes, just like in liver tissues from these animals, pathways that perform normal protein breakdown were being under-expressed which could indicate disruption to normal renal physiology (62).

**Fig 4.**
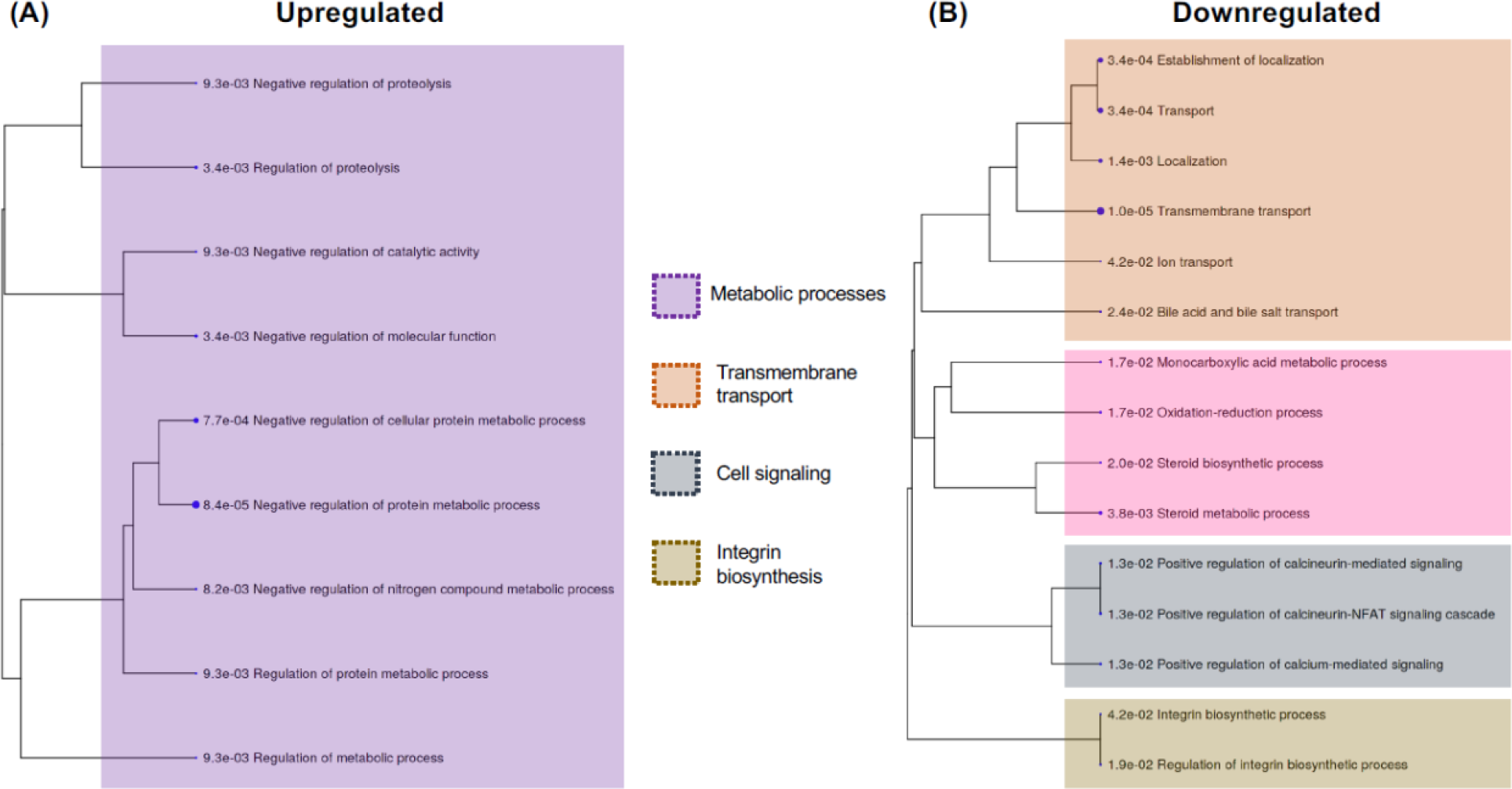
Biological processes that were differentially regulated in kidney due to SFD exposure. Hierarchical clustering tree summarizing the correlation among significant Biological process gene ontology terms (GO:BP) for genes that were (A) upregulated, or (B) downregulated in kidney. Numbers indicate p-values after FDR correction for multiple testing. Each process terms were categorized as subclass of specific parent terms as represented by colored boxes

Among the downregulated genes in kidneys of infected snakes, most belonged to biological pathways that regulate steroid metabolism, localization and transmembrane transport, and integrin synthesis (Fig. 4B). Just as in liver, steroid synthesis was also downregulated in kidney, which affects many cell-signaling pathways including electrolyte balance maintained by the kidney (63). Calcium ion signaling and transport regulates the levels of calcium in the urine and transport of bile acid and bile salts in kidney. Bile salts are important for degradation of fats, which helps in digestion and absorption of important vitamins, and elimination of toxins.

Reduced integrin activity in kidney has been linked to diseases like congenital nephrotic syndromes (CNS) and severe edema (64). Seven gene network modules were significantly associated with infection status (Table S3) and were enriched in Ribosome and N-Glycan biosynthesis KEGG pathways (Fig. S8). Lower expression of ribosome biosynthesis genes (Fig. S7) also lowers protein synthesis and causes muscle atrophy, impaired growth of new muscle fibers and loss of kidney function (62).

Taken together, our DGE and functional enrichment analysis in kidney indicate that *O. ophidiicola* infections are associated with reduced kidney function, nephrotoxicity, loss of protein metabolism, and muscular weakness. As bile is synthesized in the liver, it is likely that infection-induced liver inflammation disturbs renal functioning as well via bile salt transport pathways (65). These physiological changes are signatures of chronic kidney infections like acute kidney injury and possibly indicates that chronic *O. ophidiicola* infection induces a cascade of mechanisms that lead to multiple organ failure and eventual mortality in the host (66).

### SFD exposure triggers immune response and DNA damage repair pathways in skin

In contrast to our findings in liver and kidney, we found little evidence for differential gene expression in skin tissues between infected and control individuals (Fig.2). The overall gene expression was also lower in skin (Table S2), partly because most of the snakeskin lesions are necrotic (29) and most of the transcripts isolated from the skin of infected snakes had fungal origins (Fig. S2; Table S2). Many of the upregulated genes were part of the immune response and stress response pathways, including *JCHAIN,* which links immunoglobulin antibodies or, *PSMB8* which is likely involved in the inflammatory response pathway (Table S4). *IGLV5* encodes the variable domain of the immunoglobulin light chains that participates in the antigen recognition. Transcription factors were also upregulated in the skin, such as *ATF3,* which binds to many genomic regions that contain genes involved in cellular stress responses.

Among the genes that were downregulated (Table S4), *WNT10B* gene is expressed in epidermal keratinocytes and plays crucial roles in regulating skin development and homeostasis (67). Additionally, downregulation of *PLGC2* is associated with immune system disorders (68) and *SYP* gene, which is linked to Ca^2+^-dependent neurotransmitter release, was also under expressed in skin. Variation in SY-like immunoreactivity is a marker for neuroendocrine cancers of the skin (69).

In skin, three modules were significantly correlated with infection status and contained genes involved in DNA damage repair, cell cycle regulation, and their associated metabolic pathways like nucleic acid metabolism (Fig. S9). The modules were enriched in DNA replication and DNA mismatch repair KEGG pathways (Figs. S10, S11). Expressed in the nucleus during DNA synthesis and mismatch repair, these pathways are essential mechanisms during cell division and proliferation. Dysregulation of these pathways lead to an increase in genome instability and is associated with human nonmelanoma skin cancer (70).

Our DEG analysis of skin tissues points towards the activation of the host immune response due to infection-induced stress and rapid skin regeneration through upregulation of DNA replication and DNA repair mechanisms.

### Lower temperature exacerbates chronic infection in internal organs

Lastly, we wanted to identify if infected snakes at different temperatures showed different patterns of gene expression. Since we identified no DEGs due to interaction in skin (Fig. 2C), we focused on liver and kidney tissues. A total of 91 genes were differentially expressed in liver of infected snakes due to higher temperature, of which 41 were upregulated (45%) and 50 were downregulated (55%; Fig.5A). Functional enrichment analysis showed fibrinolysis pathways were upregulated in infected snakes at higher temperature (Fig. 5B).

**Fig 5.**
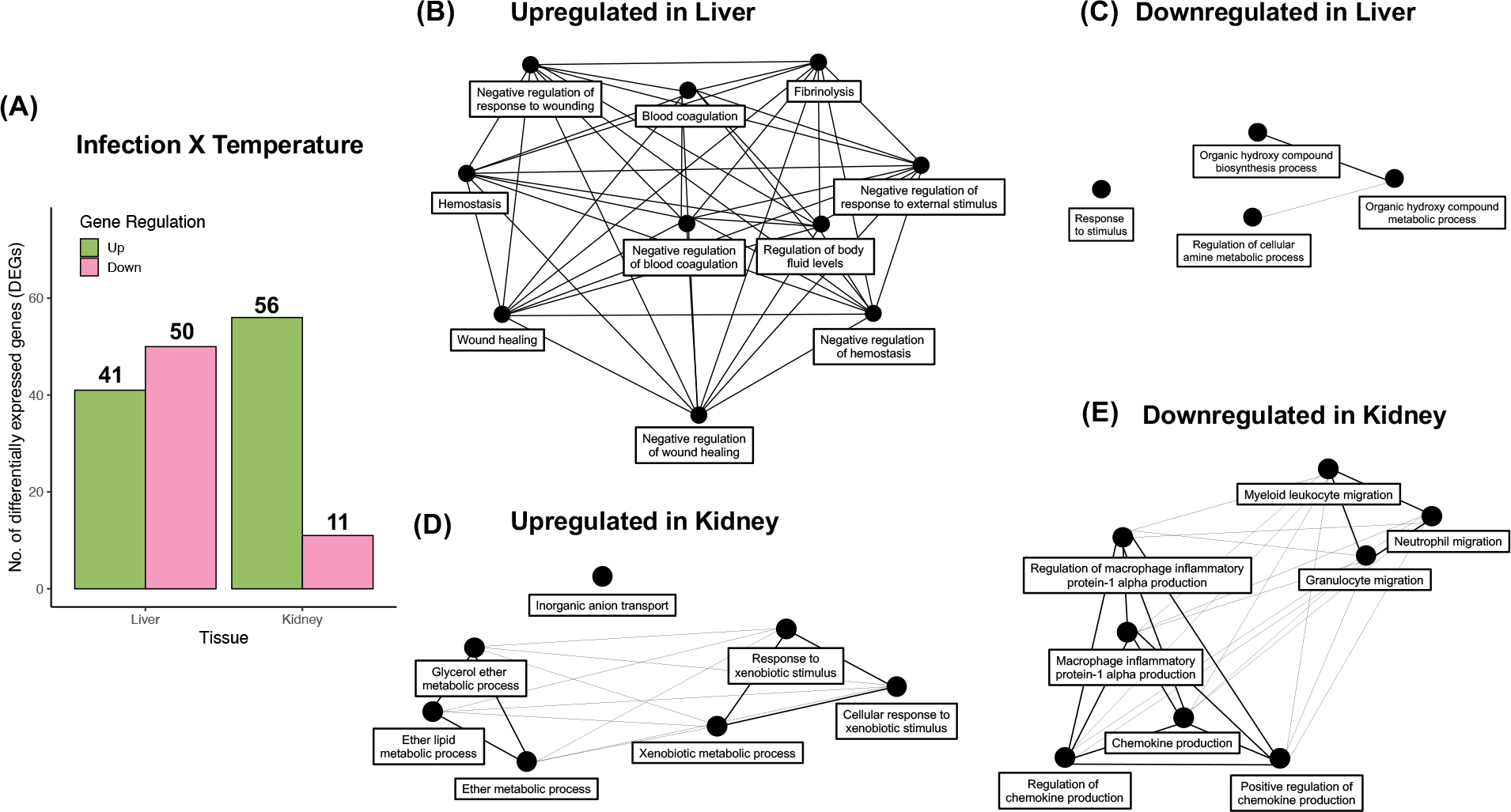
Differential expression due to interaction effect of higher temperature on infected snakes. (A) Differentially expressed genes (DEGs) in liver and kidney due to interaction between infection and higher temperature. Number above bars represent number of upregulated (green) and downregulated (pink) genes in each tissue. (B-E) Biological pathway networks for genes that were (B) upregulated in liver, (C) downregulated in liver, (D) upregulated in kidney, and (E) downregulated in kidney of infected snakes at the higher temperature as compared to infected snakes at the lower temperature. Biological pathway networks show relationship between enriched pathways (black nodes) connected by edges based on percentage of genes shared between a pair of GO:BP terms. Thicker edges represent more overlap of genes between a pair of GO term.

Fibrinolysis is the enzymatic breakdown of fibrins in blood clots. This indicates that at the lower temperature, blood coagulation processes do not stop in the livers of infected snakes, which could potentially lead to fibrosis. Most of the downregulated processes in liver due to interaction were organic hydroxy-compounds metabolic processes, possibly playing a role in alcohol metabolism (Fig. 5C) and thus, adding to already lowered protein metabolism due to SFD (Fig. 3).

In kidney tissues, we identified 67 DEGs due to interaction between treatment and temperature, of which 56 genes (84%) were upregulated and 11 (16%) genes were downregulated (Fig. 5A). Upregulated genes in infected snakes at the higher temperature were involved in response to xenobiotics and metabolism of ether lipids (Fig. 5D). Ether phospholipids metabolism is a signature of the return of blood flow to a particular organ and leads to reduction in cell death following acute kidney injury (71). Pro-inflammatory responses like production of macrophage inflammatory proteins and chemokines, and neutrophil and myeloid leukocyte migration were downregulated in infected snakes at the higher temperature (Fig. 5E). This means that infected snakes at the higher temperature showed possible signs of recovery with blood recirculation and reduced inflammatory responses.

Temperature played a crucial role in overall host response to *O. ophidiicola* infection (Fig.1) as all the infected snakes at the lower temperature were euthanized due to more severe clinical signs, whereas, most infected snakes at the higher temperature survived the experiment (Table S1). Our DGE analysis to assess the effect of temperature on gene expression suggest that infected snakes at the lower temperature were more susceptible to mortality because of multiple organ failure due to chronic infections. A negative feedback loop of higher blood coagulation and unchecked wound-healing can lead to permanent scarring and liver failure, while, prolonged kidney inflammation can induce irreversible tubular injury and nephron failure.

### Identification of potential SFD host response loci

Differential expression not only identifies a gene’s involvement in host response but also the potential contribution of products of that gene to disease resistance or susceptibility (e.g., immune genes). With the goals of characterizing potential SFD response loci in snakes we identified a total of 906 genes that were differentially regulated in at least one tissue we analyzed either due to *O. ophidiicola* infection only or due to the interaction between infection and temperature (Data S2). Within the 906 DEGs, there are 7700 exons (total length = 1,929,670 bp) in the *C. viridis* reference genome. We found 2,354 non-synonymous SNPs within the exons of DEGs that were polymorphic within the set of *C.viridis* genomes and all analyzed genomes had high diversity within these functional loci (mean heterozygosity = 0.187; SD = 0.032; Fig. S12). Though gene regulation is mostly controlled by transcription factors, sequence variation within protein coding regions is also often associated with disease susceptibility (14, 72) partly due to complex molecular co-evolution of host-pathogen system (73). Thus, these functional variants could underpin variability in host response and thus carry utility as diagnostic markers for SFD susceptibility in wild snake populations and species.

## Discussion

### Insights into SFD Host Pathology from Gene Expression Data

In terms of pathology, all wildlife fungal diseases studied to date have skin lesions as a characteristic symptom of infection (3, 6, 29) and are most likely transmitted via contact with other infected individuals (9) and/or contaminated environmental components such as soil (29) or water (6). However, despite these fungi infecting host skin epithelial cells, host response is different depending on the specific disease. For example, multiple studies involving *Bd* have confirmed an acute immune response in skin (74–76) and hinted that the early host response might be more beneficial for resistance to *Bd* infection (76). On the contrary, *P. destructans* infections on nose, wing and tail tissues in bats causes mortality as a result of a cascade of largely internal physiological changes including skin damage, dehydration, energy depletion, and higher CO2 levels in blood (77).

Our gene expression results from multiple tissues indicate that *O. ophidiicola* infection causes chronic inflammation in internal organs which damages tissues over time and disrupts normal organ physiology like protein metabolism and electrolyte balance. Our study corroborates previous expectations of SFD being a systemic disease (29) with chronic *O. ophidiicola* infections, as opposed to direct fungal damage on infected skin (41). This argues that *O. ophidiicola* pathogenesis is more similar to *P. destructans* infections in white nose syndrome where long-term exposure leads to chronic respiratory acidosis, and *O. ophidiicola* may infect the host in multiple stages to cause multiple organ failure and eventual mortality (77). We argue that a better understanding of host-pathogen interaction on snake tissues using in-vitro (e.g.(78, 79)) or in-vivo techniques (e.g. (75)) might reveal more information on the complete model of *O. ophidiicola* pathogenesis within a snake host.

More specifically, the gene expression data from skin tissue demonstrated certain similarities and differences between *Bd*, *P. destructans,* and *O. ophidiicola* infections. First, we found *O. ophidiicola* gene expression exclusively on skin (Fig. S2) and identified the expression of genes involved in hyphal growth. Necrosis on skin cell due to penetration of fungal hyphal into the epidermis have been confirmed in many SFD studies (29, 30, 80) and is the mode of *P. destructans* pathogenesis in white nose syndrome (9). Identifying functional genes within *O. ophidiicola* genome and quantifying their expression on snakeskin would help researchers identify pathogenesis mechanisms and potential deterrents against the pathogen. Comparative analysis of functional genes among different *O. ophidiicola* strains might also highlight the fungal origins and differences in virulence among lineages (7).

Second, we found the least amount of differential gene expression between infected and control snakes in skin tissue (Fig. 2). This was unexpected when compared to *Bd* as differential skin response is a major factor in determining host susceptibility to *Bd* (13) in many amphibian species (14, 18). This difference may be due to the fact that skin plays a major role in amphibian biology as it regulates osmosis and gas exchange which gets compromised and leads to death from cardiac arrest (8). Much of *O. ophidiicola* infected skin is necrotic (30) possibly terminating the gene expression of host cells. Lastly, we found over-expression of immune response and DNA damage repair genes in the skin tissues of infected snakes, which matches pro-inflammatory and skin integrity maintenance processes described in *Bd* and white nose syndrome studies (reviewed in (9, 13)). Snakes with SFD have been observed to shed more frequently (30, 81) possibly as a defense mechanism to reduce fungal load. We propose that SFD susceptibility could vary with host skin conditions and that inherent differences in snake skin (like the microdiversity) among different species/populations (81) might play a crucial role in identifying vulnerable populations for conservation. Future research should consider comparing host skin gene expression profiles in response to SFD across multiple snake species.

SFD studies in free ranging and captive snakes have documented behavioral and physiological changes in infected snakes such as lethargy (80) and higher metabolic rates (82). Much of SFD-related mortality is attributed to less frequent foraging and anorexia (30).

Starvation and anorexia cause liver injury with elevation of liver enzymes (83) and in agreement with the pro-inflammatory responses and higher metabolism we identified in liver tissues of *O. ophidiicola* infected snakes (Fig. 3). Similarly, within kidney tissues from infected snakes, reduced gene expression leads to electrolyte imbalance and loss of protein metabolism, potentially causing muscle atrophy and protein loss, and lack of liver-mediated toxin filtration (Fig. 4). Weak muscles, diseased liver, and renal failure result in lethargy in hosts and, in cases with chronic inflammation, could lead to increased impact of the disease and a higher risk of mortality.

Finally, SFD is difficult to diagnose in the wild (42) and our results suggest that possibly like as is the case of white nose syndrome, physiological impacts of SFD may start to cause deterioration of internal organs before clinical signs are manifested on the skin (77). A potential avenue for future research would be to study gene expression through time during early or late stages of *O. ophidiicola* infection to document changes before and after the onset of clinical signs. A time-course transcriptome analysis would indicate what biological changes occur when disease progresses, what genes are active at various stages of clinical signs, and what genetic and biological mechanisms underly host recovery. This can also help early diagnosis of infected snakes and identify possible genetic targets that determine host susceptibility to SFD.

### Effects of temperature

Environmental temperature plays a crucial role in fungal pathogenesis and processes like reproduction, behavior, and immune response are temperature dependent in ectotherms (13).

Many fungal species are most active at a specific range of temperatures (47) such as between 17- 25⁰C for *Bd* (84) and thus, seasonality and environmental temperature are predictors of severity in this disease (48, 85, 86). Frogs prefer warmer temperatures or induce behavioral fever to increase immune response and survival during infection (87). Snakes also display behavioral thermogenesis through basking, which increases their body temperature to increase the immune response to infection (88, 89). Our results provide a link between these behaviors and a potential response to SFD infections in snakes. Warmer environmental temperatures raise the internal body temperature beyond the critical thermal maximum of the pathogenic fungi (90) and activate immune functions like leukocyte mobility and lymphocyte response to antigens in reptiles (91). Basking has been seen reported in free ranging snakes affected by SFD (44) and is possibly a defense mechanism against infection (29). Even in our controlled experiments, all infected snakes at the lower temperature displayed more severe disease, resulting in early euthanasia (Table S1) and carried the most unique gene expression profile (Fig. 1B-D). Our results suggest that more severe disease at the lower temperature was potentially due to poor immune response, resulting in liver fibrosis and acute kidney injury in chronically inflamed tissues. Infected snakes at the higher temperature may have recovered, as our results show gene expression profile similar to control snakes at the end of the study (Fig.1B-D) including upregulation of genes that are associated with blood recirculation post-inflammation.

These results suggest that higher temperature is more suitable for recovery from SFD, but we acknowledge that we only analyzed expression profiles at a single timepoint after a significant period of exposure to the disease. Hence, it is still unclear from this study whether the lack of fungal infection and no signatures of chronic tissue injury at the higher temperature is due to recovery or lack of severe infection through the course of the experiment. An experiment that quantifies the fungal transcripts (i.e. fungal gene expression) over time would indicate whether fungus proliferates as disease progresses and declines as host recovers, or if in some cases, the fungal growth is insufficient for severe disease development and hosts clear infection before before severe clinical signs increase the likelihood of mortality.

### Potential molecular markers for diagnosing SFD susceptibility

Our inference about the physiological processes and biological pathways from differentially expressed genes was based on functions predicted from orthologs. A caveat with this approach is that GO terms are only available for small set of model organisms and functional impact of many species-specific genes not identified in the genome annotation of the model organism are unknown. Relying on functional information of homologous genes in related species (*Anolis carolinensis* in this study) does yield a comprehensive inventory of genes and their functional involvement in disease response.. For example, skin integrity genes are upregulated in *Bd*-resistant species (18, 74) and downregulated in *Bd*-susceptible species (74) across multiple studies (13).

*O. ophidiicola* is a generalist pathogen which is capable of infecting numerous snake species across the phylogeny (31, 33, 92). We acknowledge that the physiological response to this pathogen may vary among different host snake species, but we suspect that the biological mechanisms underlying host response (like immune activation, metabolism, etc.) would be similar among different snake species. Our curated set of genes (Data S1-2) can be used to compare vulnerability to SFD in different species, specifically how changes in gene expression and/or certain polymorphisms are associated with SFD response. For example, the link between lower acquired immune gene expression and *Bd* survival has been observed in many amphibian species (14, 93). and specific genotypes can be associated with disease resistance (14, 72).

Differences in allele specific variation in expression (94) or genotype association due to host-pathogen co-evolution like MHC allelic associations observed in case of amphibians exposed to *Bd* (14, 72) or natural populations of bananaquits with chronic avian malaria (95) can be leveraged to assess genetic vulnerability to pathogenic diseases. *O. ophiodiicola* has likely been prevailing in nature longer than previously expected (36) and thus, certain snake species could possibly have evolved to resist fungal infection due to long-term pathogenic pressure. Thus, comparing variation in sequence diversity and/or expression profiles of candidate genes among host populations and species host can help conservation biologist provide diagnostic capability to identify vulnerable populations most likely to succumb to SFD.

## Conclusion

Ophidiomycosis (snake fungal disease; SFD) is a recently identified fungal disease and a potential threat to ecologically and phylogenetically diverse snake species. Our transcriptomic results demonstrate that similar to white nose syndrome in bats and in contrast to chytridiomycosis in amphibians, SFD is also a systemic disease. Fungal infections were localized to the skin, as expected, but most physiological changes occurred due to chronic inflammation in the internal organs. Temperature played an important role as infected snakes maintained at the lower temperature had higher mortality and a unique gene expression profile; this suggests that higher temperatures possibly aid in recovery from the disease. Future research should use multi-omic approaches to determine whether different *O. ophiodiicola* strains produce similar impacts among snakes and whether different snake species and populations vary in their response to SFD. Finally, our study reports a set of candidate genes and functional loci that carry significance in improving our understanding of host-pathogen dynamics in multiple species and can help diagnose and mitigate SFD in wild and captive snakes.

## Materials and Methods

### Controlled experiment varying infection status and temperature

Twelve adult prairie rattlesnakes (*Crotalus viridis*) were evenly divided and maintained at 20°C or 26°C following a two-week acclimatization period. These temperatures were chosen as possible ends of the *O. ophidiicola* thermal optimum ((31); thermal optimum for *Bd* is 17-25⁰C (81)). Four animals at each temperature condition were randomly selected for experimental challenge while two animals were maintained as controls (Fig. 1A). All snakes were free of clinical signs before inoculation based on criteria previously described (96). All snakes were offered pre-killed mice every 1–2 weeks and water was provided *ad libitum*. A pure *O. ophidiicola* isolate (UI-VDL # 12–34933) cultured from an eastern massasauga (*Sistrurus catenatus*) was injected intradermally (0.1 ml of a pure culture containing 109,000 CFUs). The control animals were inoculated with a similar volume of sterile saline. All injections were performed over the dorsal mid-body of each snake. The experiment was conducted for 90 days post infection (dpi) and snakes were euthanized at the end of the study or earlier in case of severe disease mortality by complete anesthesia with ketamine intramuscularly, followed by sodium pentobarbital intravenously. All procedures were approved by the University of Illinois Institutional Animal Care and Use Committee (IACUC; Protocol: 19165). However, during the experiment, one of the infected snakes at the higher temperature had to be prematurely euthanized before tissues could be extracted. Each animal was subjected to necropsy examination separately to collect samples of liver, kidney and skin. RNA was extracted using Qiagen RNEasy kit according to the manufacturer’s recommendations (Qiagen RNEasy, Valencia, CA). RNA samples were analyzed using spectrophotometry (NanoDrop 1000, Thermo Fisher) to quantify concentration and purity, then stored at -80°C. Reverse transcription was performed using the Quantitect Reverse Transcription kit (Qiagen, Valencia, CA) following manufacturer’s protocol. The cDNA was submitted to the Keck Biotechnology Center at the University of Illinois for further processing.

### Construction and sequencing of RNAseq libraries

RNAseq libraries were prepared for each sample with Illumina’s TruSeq Stranded mRNAseq Sample Prep kit and paired-end (PE) reads (2x150bp) were sequenced on SP-lane of Illumina NovaSeq 6000 sequencer at the University of Illinois Roy Carver Genome Center.

Following sequencing, we removed 1 skin and 1 liver sample due to low number of raw sequences generated. Ultimately, we analyzed sequence data from 10 liver, 11 kidney, and 10 skin tissues from each snake with different temperature condition and infection status (Fig. 1A; Table S1).

### Sequence processing and expression quantification

We removed adapter sequences and clipped poor-quality bases (quality score < 20) from both ends of raw reads from the sequencer using Trimmomatic (97), aligned filtered reads to the annotated *C. viridis* reference genome (UTA_CroVir_3.0; GenBank assembly accession: GCA_003400415.2) using hisat2 (98) with the *–downstream-transcriptome-assembly* option and reporting primary alignments. We next assembled transcripts for each sample using StringTie (99) default parameters and *C. viridis* reference annotation file to guide assembly and merged sample transcripts using StringTie. Next, a transcript count matrix was next created with featureCounts (100) which excluded chimeric fragments and only retained mapped fragments in which both read pairs successfully aligned to the reference. We only counted reads that matched to the exons present in the merged annotation file. Ultimately, we generated a table of the count data which reports, for each sample, the number of sequence fragments that were assigned to each transcript.

To identify the number of total transcripts from snake tissues that had fungal origins, we mapped the aligned filtered reads to *O. ophidiicola* reference genome (GenBank assembly accession GCA_002167195.1) using hisat2. We expect that tissues where fungus is actively present would have more expression of fungal transcripts and thus would have higher alignment rate to the *O. ophidiicola* reference genome.

### Differential gene expression analysis

We conducted differential gene expression (DGE) analyses from the counts of all assembled transcripts (N = 107,167) for each tissue separately using DESeq2 (101) in R (102) following the workflow as described by the authors (101, 103). Our statistical model used a multi-factor design that included both main factors and an interaction term. The main factors were temperature (“Temperature”; low [20°C] vs. high [26°C]) and infection status (“Infection”; *O. ophidiicola* infected vs. control) and an interaction terms that measures an interaction effect between temperature and infection (Infection X Temperature) (Fig. 1A). By analyzing genes showing differential gene expression due to each of the main factors and the interaction term, we were able to identify genes with differential expression purely due to the effect of treatment, temperature, or the interaction between these factors (Fig. S4).

To quantify if there was any difference in gene expression between treated tissues due to differences in temperatures, we used *contrast* function in DESeq2. To remove potential false positives due to low expression, we applied independent filtering to remove transcripts with low read counts and only considered transcripts with a false discovery rate (FDR) adjusted p-value of <0.05 to be differentially expressed. Finally, we transformed log fold change data using “adaptive shrinkage” from *ashr* package (104) in R to shrink the change in expression to only retain differentially expressed genes (DEGs) that are most biologically significant.

### Weighted gene co-expression network analysis

Besides identifying individual genes that showed differential expression in each tissue due to differences in treatment and temperature conditions, we also identified networks of differentially co-expressed genes (termed “modules”) from the normalized counts of all expressed genes (105). Estimating co-expression patterns can provide insights into the biological processes that underlie the complex cascade of events that lead to the phenotypic differences observed due to different treatment effects (106). The nodes of these co-expression gene networks correspond to gene expression profiles, and edges between genes are determined by the pairwise correlations between the level of expression of each gene (105).

We used weighted gene co-expression network analysis (WGCNA) (107) to cluster highly correlated genes into different modules and to then quantify the relationship of different modules with each other and to the temperature and infection treatments following the workflow described by the authors (https://horvath.genetics.ucla.edu/html/CoexpressionNetwork/Rpackages/WGCNA/Tutorials/index.html;last accessed 03/15/2022). We first identified modules of co-expressed genes by calculating pairwise Pearson correlations between each pair of genes with non-zero expression in at least 8 of our samples in each tissue separately (Ngenes_liver = 45,021; Ngenes_kidney = 47,522; Ngenes_skin = 44,807). We then merged modules that were correlated to each other with R^2^ > 0.75 to get a final set of merged modules. Modules were identified by hierarchical clustering of signed Topological Overlap Matrix (TOM) (108) using a soft threshold (β) to assign a connection weight to each gene pair and minimum 30 co-expressed genes to be assigned to a module following (109). β was chosen as the lowest power for which the scale-free topology fit index of all genes reached 0.9 within each tissue type (βliver = 12; βkidney = 8; βskin = 12) as suggested by the authors of WGCNA.

Next, we identified modules that were significantly associated (P < 0.05 after applying false discovery rate method to correct for multiple testing) with the external temperature and infection treatments. We only retained modules that were significantly correlated to (a) either infection or (b) both infection and temperature. Within each significant module, we also identified the module membership and gene significance for each gene within the module.

Module membership measures correlation between a gene’s expression profile with the module eigengene of a given module. Highly connected genes within a module are more likely to have higher module membership values to the respective module. Gene significance is the measure of correlation of a gene with an external treatment factor (infection and temperature) and indicates the biological significance of a module gene with respect to the fixed effects of our experimental design. For many of our final modules, genes with higher module membership had a positive and significant correlation with gene significance meaning that highly connected genes within a module are more likely to be associated with experimental treatments. Finally, we matched the genes belonging to final set of modules and were also differentially expressed due to the infection treatment only.

### Functional enrichment analysis

For each module that was significantly associated with treatment each tissue and DEGs, we performed a functional enrichment analysis to identify which biological process gene ontology (GO:BP) terms and KEGG pathways were overrepresented in those gene clusters, as compared to the genome-wide GO complements of *Anolis carolinensis*. We performed GO enrichment analysis for all DEGs that were upregulated or downregulated within each tissue separately. We first converted candidate annotated genes in *C. viridis* genome to *A. carolinensis* Ensembl IDs using DAVID (110, 111) and performed functional enrichment analysis using gProfiler (112). We used g:SCS method (significance threshold < 0.05) for computing multiple testing correction for p-values gained from GO and pathway enrichment analysis to account for non-independence among multiple tests among GO terms (112). We then created phylogenetic trees and functional networks using ShinyGO (113) to visualize enriched pathways.

### Identification of potential SFD host response loci

Finally, to identify potential SFD host response loci that could be used to assay possible disease susceptibility or resistance at the population and species level in snakes, we first combined all the DEGs within all the tissues (Data S2) and then extracted protein coding regions

(exons) within the DEGs using the *C. viridis* genome annotation (see above). Next, to look for non-synonymous mutations within DEGs that could be potential diagnostic markers, we first mapped 19 *C. viridis* genomes (BioProject No. PRJNA593834; SRA Nos. SRS5767847-59, SRS5767870, SRS5767880-85) to the reference genome and then, identified single nucleotide polymorphism (SNP) markers within DEG exons (see Supplementary Text S1 for details).

### Data Availability Statement

The sequence datasets generated during the current study are available in NCBI’s Short Read Archive BioProject Accession No. PRJNA817280, BioSample Accession Nos. SAMN26754183-4213 and SRA Accession Nos. SRR18361039-069. The scripts developed for analysis can be publicly accessed at https://github.com/samarth8392/SFDTranscriptomics.

## Acknowledgements

This work was supported by the State Wildlife Grants Program, administered jointly by the US Fish and Wildlife Service and the Ohio Division of Wildlife, with funds provided by the Ohio Biodiversity Conservation Partnership between Ohio State University and the Ohio Division of Wildlife. HLG was also supported by National Science Foundation (USA) Grant DEB 1638872. We thank Allison Wright, Michelle Waligora, Kennymac Durante, and Kelcie Fredrickson for care of the rattlesnakes throughout the study. We thank Alvaro Hernandez and his team at Roy Carver Genomics Center at University of Illinois, Urbana Champaign for RNA sequencing services.

## Author Contributions

**Conceptualization**: Samarth Mathur, Ellen Haynes, Matthew C. Allender, H. Lisle Gibbs

**Data curation**: Samarth Mathur, H. Lisle Gibbs

**Analysis**: Samarth Mathur

**Funding acquisition**: Matthew C. Allender, H. Lisle Gibbs

**Investigation**: Samarth Mathur, H. Lisle Gibbs

**Methodology**: Samarth Mathur, H. Lisle Gibbs

**Project administration**: H. Lisle Gibbs, Matthew C. Allender

**Resources**: H. Lisle Gibbs, Matthew C. Allender

**Visualization**: Samarth Mathur, H. Lisle Gibbs

**Writing – original draft:** Samarth Mathur, H. Lisle Gibbs.

**Writing – review & editing**: Samarth Mathur, Ellen Haynes, Matthew C. Allender, H. Lisle Gibbs

## References

1. Becker K, Hu Y, Biller-Andorno N. Infectious diseases – A global challenge. International Journal of Medical Microbiology. 2006;296(4-5):179–85.

2. Barroso P, Acevedo P, Vicente J. The importance of long-term studies on wildlife diseases and their interfaces with humans and domestic animals: A review. Transboundary and Emerging Diseases. 2020;68(4):1895–909.

3. Berger L, Speare R, Daszak P, Green DE, Cunningham AA, Goggin CL, et al. Chytridiomycosis causes amphibian mortality associated with population declines in the rain forests of Australia and Central America. Proceedings of the National Academy of Sciences. 1998;95(15):9031–6.

4. Frick WF, Pollock JF, Hicks AC, Langwig KE, Reynolds DS, Turner GG, et al. An emerging disease causes regional population collapse of a common North American bat species. Science. 2010;329(5992):679-82.

5. Fisher MC, Garner TWJ. Chytrid fungi and global amphibian declines. Nat Rev Microbiol. 2020;18(6):332–43.

6. Kilpatrick AM, Briggs CJ, Daszak P. The ecology and impact of chytridiomycosis: an emerging disease of amphibians. Trends in Ecology & Evolution. 2010;25(2):109–18.

7. Rosenblum EB, James TY, Zamudio KR, Poorten TJ, Ilut D, Rodriguez D, et al. Complex history of the amphibian-killing chytrid fungus revealed with genome resequencing data. Proceedings of the National Academy of Sciences. 2013;110(23):9385–90.

8. Voyles J, Young S, Berger L, Campbell C, Voyles WF, Dinudom A, et al. Pathogenesis of Chytridiomycosis, a Cause of Catastrophic Amphibian Declines. Science. 2009;326(5952):582–5.

9. Hoyt JR, Kilpatrick AM, Langwig KE. Ecology and impacts of white-nose syndrome on bats. Nature Reviews Microbiology. 2021;19(3):196–210.

10. Frick WF, Puechmaille SJ, Hoyt JR, Nickel BA, Langwig KE, Foster JT, et al. Disease alters macroecological patterns of North American bats. Global Ecology and Biogeography. 2015;24(7):741–9.

11. Meteyer CU, Buckles EL, Blehert DS, Hicks AC, Green DE, Shearn-Bochsler V, et al. Histopathologic criteria to confirm white-nose syndrome in bats. Journal of Veterinary Diagnostic Investigation. 2009;21(4):411–4.

12. Arlettaz R, Reeder DM, Frank CL, Turner GG, Meteyer CU, Kurta A, et al. Frequent arousal from hibernation linked to severity of infection and mortality in bats with white-nose syndrome. PLoS ONE. 2012;7(6).

13. Zamudio KR, McDonald CA, Belasen AM. High variability in infection mechanisms and host responses: a review of functional genomic studies of amphibian chytridiomycosis. Herpetologica. 2020;76(2).

14. McDonald CA, Longo AV, Lips KR, Zamudio KR. Incapacitating effects of fungal coinfection in a novel pathogen system. Mol Ecol. 2020;29(17):3173–86.

15. Andre SE, Parker J, Briggs CJ. Effect of temperature on host response to *Batrachochytrium dendrobatidis* infection in the mountain yellow-legged frog (*Rana muscosa*). Journal of Wildlife Diseases. 2008;44(3):716–20.

16. Brannelly LA, Berger L, Marrantelli G, Skerratt LF. Low humidity is a failed treatment option for chytridiomycosis in the critically endangered southern corroboree frog. Wildlife Research. 2015;42(1).

17. Grogan LF, Robert J, Berger L, Skerratt LF, Scheele BC, Castley JG, et al. Review of the amphibian immune response to chytridiomycosis, and future directions. Frontiers in Immunology. 2018;9.

18. Eskew EA, Shock BC, LaDouceur EEB, Keel K, Miller MR, Foley JE, et al. Gene expression differs in susceptible and resistant amphibians exposed to *Batrachochytrium dendrobatidis*. Royal Society Open Science. 2018;5(2):170910.

19. Sarmiento-Ramírez JM, Abella E, Martín MP, Tellería MT, López-Jurado LF, Marco A, et al. *Fusarium solani* is responsible for mass mortalities in nests of loggerhead sea turtle, *Caretta caretta*, in Boavista, Cape Verde. FEMS Microbiology Letters. 2010;312(2):192–200.

20. Kim K, Harvell CD. The rise and fall of a six-year coral-fungal epizootic. The American Naturalist. 2004;164(S5):S52–S63.

21. Schumacher J. Fungal diseases of reptiles. Veterinary Clinics of North America: Exotic Animal Practice. 2003;6(2):327–35.

22. Brown J, vanEngelsdorp D, Evans JD, Saegerman C, Mullin C, Haubruge E, et al. Colony collapse disorder: a descriptive study. PLoS ONE. 2009;4(8).

23. Anderson PK, Cunningham AA, Patel NG, Morales FJ, Epstein PR, Daszak P. Emerging infectious diseases of plants: pathogen pollution, climate change and agrotechnology drivers. Trends in Ecology & Evolution. 2004;19(10):535–44.

24. Brown JKM, Hovmøller MS. Aerial dispersal of pathogens on the global and continental scales and its impact on plant disease. Science. 2002;297(5581):537–41.

25. Fisher MC, Henk DA, Briggs CJ, Brownstein JS, Madoff LC, McCraw SL, et al. Emerging fungal threats to animal, plant and ecosystem health. Nature. 2012;484(7393):186–94.

26. Ghosh PN, Fisher MC, Bates KA. Diagnosing emerging fungal threats: A one health perspective. Frontiers in Genetics. 2018;9.

27. Williams E, Yuill T, Artois M, Fischer J, Haigh S. Emerging infectious diseases in wildlife. Revue scientifique et technique-Office international des Epizooties. 2002;21(1):139–58.

28. Daszak P, Cunningham AA, Hyatt AD. Emerging infectious diseases of wildlife-- threats to biodiversity and human health. Science. 2000;287(5452):443–9.

29. Lorch JM, Knowles S, Lankton JS, Michell K, Edwards JL, Kapfer JM, et al. Snake fungal disease: an emerging threat to wild snakes. Philos Trans R Soc Lond B Biol Sci. 2016;371(1709).

30. Lorch JM, Lankton J, Werner K, Falendysz EA, McCurley K, Blehert DS. Experimental infection of snakes with *Ophidiomyces ophiodiicola* causes pathological changes that typify snake fungal disease. mBio. 2015;6(6):e01534–15.

31. Allender MC, Raudabaugh DB, Gleason FH, Miller AN. The natural history, ecology, and epidemiology of *Ophidiomyces ophiodiicola* and its potential impact on free-ranging snake populations. Fungal Ecology. 2015;17:187–96.

32. Rajeev S, Sutton DA, Wickes BL, Miller DL, Giri D, Van Meter M, et al. Isolation and characterization of a new fungal species, *Chrysosporium ophiodiicola*, from a mycotic granuloma of a black rat snake (*Elaphe obsoleta obsoleta*). J Clin Microbiol. 2009;47(4):1264–8.

33. Burbrink FT, Lorch JM, Lips KR. Host susceptibility to snake fungal disease is highly dispersed across phylogenetic and functional trait space. Sci Adv. 2017;3(12):e1701387.

34. Allender MC, Dreslik M, Wylie S, Phillips C, Wylie DB, Maddox C, et al. *Chrysosporium* sp. Infection in eastern massasauga rattlesnakes. Emerging infectious diseases. 2011;17(12):2383–4.

35. Clark RW, Marchand MN, Clifford BJ, Stechert R, Stephens S. Decline of an isolated timber rattlesnake (*Crotalus horridus*) population: Interactions between climate change, disease, and loss of genetic diversity. Biological Conservation. 2011;144(2):886–91.

36. Lorch JM, Price SJ, Lankton JS, Drayer AN. Confirmed Cases of Ophidiomycosis in Museum Specimens from as Early as 1945, United States. Emerg Infect Dis. 2021;27(7):1986–9.

37. Sigler L, Hambleton S, Paré JA. Molecular characterization of reptile pathogens currently known as members of the chrysosporium anamorph of *Nannizziopsis vriesii* complex and relationship with some human-associated isolates. Journal of Clinical Microbiology. 2013;51(10):3338–57.

38. Sun PL, Yang CK, Li WT, Lai WY, Fan YC, Huang HC, et al. Infection with *Nannizziopsis guarroi* and *Ophidiomyces ophiodiicola* in reptiles in Taiwan. Transboundary and Emerging Diseases. 2021.

39. Takami Y, Nam KO, Takaki Y, Kadekaru S, Hemmi C, Hosoya T, et al. First report of ophidiomycosis in Asia caused by *Ophidiomyces ophiodiicola* in captive snakes in Japan. J Vet Med Sci. 2021;83(8):1234–9.

40. Nichols DK, Weyant RS, Lamirande EW, Sigler L, Mason RT. Fatal mycotic dermatitis in captive brown tree snakes (*Boiga irregularis*). J Zoo Wildl Med. 1999;30(1):111–8.

41. Dolinski AC, Allender MC, Hsiao V, Maddox CW. Systemic *Ophidiomyces ophiodiicola* infection in a free-ranging plains garter snake (*Thamnophis radix*). Journal of Herpetological Medicine and Surgery. 2014;24(1).

42. Davy CM, Shirose L, Campbell D, Dillon R, McKenzie C, Nemeth N, et al. Revisiting ophidiomycosis (snake fungal disease) after a decade of targeted research. Frontiers in Veterinary Science. 2021;8.

43. Guthrie AL, Knowles S, Ballmann AE, Lorch JM. Detection of snake fungal disease due to *Ophidiomyces ophiodiicola* in virginia, usa. Journal of Wildlife Diseases. 2016;52(1):143–9.

44. McBride MP, Wojick KB, Georoff TA, Kimbro J, Garner MM, Wang X, et al. *Ophidiomyces ophiodiicola* dermatitis in eight free-ranging timber rattlesnakes (*Crotalus horridus*) from Massachusetts. Journal of Zoo and Wildlife Medicine. 2015;46(1):86–94.

45. McCoy CM, Lind CM, Farrell TM. Environmental and physiological correlates of the severity of clinical signs of snake fungal disease in a population of pigmy rattlesnakes, Sistrurus miliarius. Conservation Physiology. 2017;5(1).

46. Lind CM, McCoy CM, Farrell TM. Tracking outcomes of snake fungal disease in free-ranging pygmy rattlesnakes (*Sistrurus miliarius*). Journal of Wildlife Diseases. 2018;54(2):352–6.

47. Robert Vincent A, Casadevall A. Vertebrate endothermy restricts most fungi as potential pathogens. The Journal of Infectious Diseases. 2009;200(10):1623–6.

48. Rowley JJL, Alford RA. Hot bodies protect amphibians against chytrid infection in nature. Scientific Reports. 2013;3(1).

49. Ghosh PN, Brookes LM, Edwards HM, Fisher MC, Jervis P, Kappel D, et al. Cross-disciplinary genomics approaches to studying emerging fungal infections. Life. 2020;10(12).

50. Andrianopoulos A, Tang G, Chen Y, Xu J-R, Kistler HC, Ma Z. The fungal myosin I is essential for Fusarium toxisome formation. PLOS Pathogens. 2018;14(1).

51. Prostak SM, Robinson KA, Titus MA, Fritz-Laylin LK. The actin networks of chytrid fungi reveal evolutionary loss of cytoskeletal complexity in the fungal kingdom. Current Biology. 2021;31(6):1192–205.e6.

52. Renshaw H, Juvvadi PR, Cole DC, Steinbach WJ. The class V myosin interactome of the human pathogen *Aspergillus fumigatus* reveals novel interactions with COPII vesicle transport proteins. Biochemical and Biophysical Research Communications. 2020;527(1):232–7.

53. Huang C, Feng Y, Patel G, Xu X-q, Qian J, Liu Q, et al. Production, immobilization and characterization of beta-glucosidase for application in cellulose degradation from a novel *Aspergillus versicolor*. International Journal of Biological Macromolecules. 2021;177:437–46.

54. Burnie JP, Carter TL, Hodgetts SJ, Matthews RC. Fungal heat-shock proteins in human disease. FEMS Microbiology Reviews. 2006;30(1):53–88.

55. Sherr CJ. Colony-stimulating factor-1 receptor. Blood. 1990;75(1):1–12.

56. Qin M, Liang Z, Qin H, Huo Y, Wu Q, Yang H, et al. Novel prognostic biomarkers in gastric cancer: CGB5, MKNK2, and PAPPA2. Frontiers in Oncology. 2021;11.

57. Yang M, Ma F, Guan M. Role of steroid hormones in the pathogenesis of nonalcoholic fatty liver disease. Metabolites. 2021;11(5).

58. Scheller K, Sekeris CE. The effects of steroid hormones on the transcription of genes encoding enzymes of oxidative phosphorylation. Experimental Physiology. 2003;88(1):129–40.

59. Kersten S, Stienstra R. The role and regulation of the peroxisome proliferator activated receptor alpha in human liver. Biochimie. 2017;136:75–84.

60. Younossi ZM, Stepanova M, Afendy M, Fang Y, Younossi Y, Mir H, et al. Changes in the prevalence of the most common causes of chronic liver diseases in the United States from 1988 to 2008. Clinical Gastroenterology and Hepatology. 2011;9(6):524–30.e1.

61. Lecker SH, Mitch WE. Proteolysis by the ubiquitin-proteasome system and kidney disease. Journal of the American Society of Nephrology. 2011;22(5):821–4.

62. Wang XH, Mitch WE. Mechanisms of muscle wasting in chronic kidney disease. Nature Reviews Nephrology. 2014;10(9):504–16.

63. Lameire N, Van Biesen W, Vanholder R. Electrolyte disturbances and acute kidney injury in patients with cancer. Seminars in Nephrology. 2010;30(6):534–47.

64. Kreidberg JA, Symons JM. Integrins in kidney development, function, and disease. American Journal of Physiology-Renal Physiology. 2000;279(2):F233–F42.

65. El Chediak A, Janom K, Koubar SH. Bile cast nephropathy: when the kidneys turn yellow. Renal Replacement Therapy. 2020;6(1).

66. Schardong J, Marcolino MAZ, Plentz RDM. Muscle atrophy in chronic kidney disease. Advances in Experimental Medicine and Biology 2018. p. 393–412.

67. Ouji Y, Yoshikawa M, Shiroi A, Ishizaka S. *Wnt-10b* promotes differentiation of skin epithelial cells in vitro. Biochemical and Biophysical Research Communications. 2006;342(1):28–35.

68. Zhou Q, Lee G-S, Brady J, Datta S, Katan M, Sheikh A, et al. A hypermorphic missense mutation in *PLCG2*, encoding phospholipase Cγ2, causes a dominantly inherited autoinflammatory disease with immunodeficiency. The American Journal of Human Genetics. 2012;91(4):713–20.

69. Hamie L, Abbas O, Bhawan J. Neuroendocrine differentiation of skin tumors: a comprehensive review. The American Journal of Dermatopathology. 2020;42(12):899–910.

70. Li G-M. Mechanisms and functions of DNA mismatch repair. Cell Research. 2007;18(1):85–98.

71. Yan H-f, Zou T, Tuo Q-z, Xu S, Li H, Belaidi AA, et al. Ferroptosis: mechanisms and links with diseases. Signal Transduction and Targeted Therapy. 2021;6(1).

72. Savage AE, Zamudio KR. MHC genotypes associate with resistance to a frog-killing fungus. Proceedings of the National Academy of Sciences. 2011;108(40):16705–10.

73. Phillips KP, Cable J, Mohammed RS, Herdegen-Radwan M, Raubic J, Przesmycka KJ, et al. Immunogenetic novelty confers a selective advantage in host–pathogen coevolution. Proceedings of the National Academy of Sciences. 2018;115(7):1552–7.

74. Ellison AR, Tunstall T, DiRenzo GV, Hughey MC, Rebollar EA, Belden LK, et al. More than skin deep: functional genomic basis for resistance to amphibian chytridiomycosis. Genome Biology and Evolution. 2015;7(1):286–98.

75. Ellison AR, DiRenzo GV, McDonald CA, Lips KR, Zamudio KR. First in vivo *Batrachochytrium dendrobatidis* transcriptomes reveal mechanisms of host exploitation, host- specific gene expression, and expressed genotype shifts. G3 Genes|Genomes|Genetics. 2017;7(1):269–78.

76. Grogan LF, Cashins SD, Skerratt LF, Berger L, McFadden MS, Harlow P, et al. Evolution of resistance to chytridiomycosis is associated with a robust early immune response. Molecular Ecology. 2018;27(4):919–34.

77. Verant ML, Meteyer CU, Speakman JR, Cryan PM, Lorch JM, Blehert DS. White-nose syndrome initiates a cascade of physiologic disturbances in the hibernating bat host. BMC Physiology. 2014;14(1).

78. Fites JS, Ramsey JP, Holden WM, Collier SP, Sutherland DM, Reinert LK, et al. The invasive chytrid fungus of amphibians paralyzes lymphocyte responses. Science. 2013;342(6156):366–9.

79. Fisher MC, Rosenblum EB, Poorten TJ, Joneson S, Settles M. Substrate-specific gene expression in *Batrachochytrium dendrobatidis*, the chytrid pathogen of amphibians. PLoS ONE. 2012;7(11).

80. McKenzie CM, Oesterle PT, Stevens B, Shirose L, Mastromonaco GF, Lillie BN, et al. Ophidiomycosis in red cornsnakes (*Pantherophis guttatus*): potential roles of brumation and temperature on pathogenesis and transmission. Vet Pathol. 2020;57(6):825–37.

81. Allender MC, Baker S, Britton M, Kent AD. Snake fungal disease alters skin bacterial and fungal diversity in an endangered rattlesnake. Scientific Reports. 2018;8(1).

82. Agugliaro J, Lind CM, Lorch JM, Farrell TM, Hawley D. An emerging fungal pathogen is associated with increased resting metabolic rate and total evaporative water loss rate in a winter- active snake. Functional Ecology. 2019;34(2):486–96.

83. Rosen E, Bakshi N, Watters A, Rosen HR, Mehler PS. Hepatic complications of anorexia nervosa. Digestive Diseases and Sciences. 2017;62(11):2977–81.

84. Piotrowski JS, Annis SL, Longcore JE. Physiology of *Batrachochytrium dendrobatidis*, a chytrid pathogen of amphibians. Mycologia. 2017;96(1):9–15.

85. Longo AV, Burrowes PA, Joglar RL. Seasonality of *Batrachochytrium dendrobatidis* infection in direct-developing frogs suggests a mechanism for persistence. Diseases of Aquatic Organisms. 2009;92(3):253–60.

86. Longo AV, Zamudio KR. Temperature variation, bacterial diversity and fungal infection dynamics in the amphibian skin. Molecular Ecology. 2017;26(18):4787–97.

87. Boltaña S, Rey S, Roher N, Vargas R, Huerta M, Huntingford FA, et al. Behavioural fever is a synergic signal amplifying the innate immune response. Proceedings of the Royal Society B: Biological Sciences. 2013;280(1766).

88. Burns G, Ramos A, Muchlinski A. Fever response in North American snakes. Journal of Herpetology. 1996;30(2).

89. Godwin CD, Walker DM, Romer AS, Grajal-Puche A, Grisnik M, Goessling JM, et al. Testing the febrile response of snakes inoculated with *Ophidiomyces ophidiicola*, the causative agent of snake fungal disease. Journal of Thermal Biology. 2021;100.

90. Zimmerman LM, Vogel LA, Bowden RM. Understanding the vertebrate immune system: insights from the reptilian perspective. Journal of Experimental Biology. 2010;213(5):661–71.

91. Vaughn LK, Bernheim HA, Kluger MJ. Fever in the lizard *Dipsosaurus dorsalis*. Nature. 1974;252(5483):473–4.

92. Allender MC, Ravesi MJ, Haynes E, Ospina E, Petersen C, Phillips CA, et al. Ophidiomycosis, an emerging fungal disease of snakes: Targeted surveillance on military lands and detection in the western US and Puerto Rico. Plos One. 2020;15(10).

93. Savage AE, Gratwicke B, Hope K, Bronikowski E, Fleischer RC. Sustained immune activation is associated with susceptibility to the amphibian chytrid fungus. Mol Ecol. 2020;29(15):2889–903.

94. Knight JC. Allele-specific gene expression uncovered. Trends in Genetics. 2004;20(3):113–6.

95. Antonides J, Mathur S, Sundaram M, Ricklefs R, DeWoody JA. Immunogenetic response of the bananaquit in the face of malarial parasites. BMC Evolutionary Biology. 2019;19(1).

96. Baker SJ, Haynes E, Gramhofer M, Stanford K, Bailey S, Christman M, et al. Case definition and diagnostic testing for snake fungal disease. Herpetological Review. 2019;50(2):279–85.

97. Bolger AM, Lohse M, Usadel B. Trimmomatic: a flexible trimmer for Illumina sequence data. Bioinformatics. 2014;30(15):2114–20.

98. Kim D, Langmead B, Salzberg SL. HISAT: a fast spliced aligner with low memory requirements. Nature Methods. 2015;12(4):357–60.

99. Pertea M, Pertea GM, Antonescu CM, Chang T-C, Mendell JT, Salzberg SL. StringTie enables improved reconstruction of a transcriptome from RNA-seq reads. Nature Biotechnology. 2015;33(3):290–5.

100. Liao Y, Smyth GK, Shi W. featureCounts: an efficient general purpose program for assigning sequence reads to genomic features. Bioinformatics. 2013;30(7):923–30.

101. Love MI, Huber W, Anders S. Moderated estimation of fold change and dispersion for RNA-seq data with DESeq2. Genome Biol. 2014;15(12):550.

102. Team RC. R: A language and environment for statistical computing. 2013.

103. Love M, Anders S, Kim V, Huber W. RNA-Seq workflow: gene-level exploratory analysis and differential expression. F1000Research. 2016;4(1070).

104. Stephens M. False discovery rates: a new deal. Biostatistics. 2016.

105. Zhang B, Horvath S. A general framework for weighted gene co-expression network analysis. Statistical Applications in Genetics and Molecular Biology. 2005;4(1).

106. Nacu S, Critchley-Thorne R, Lee P, Holmes S. Gene expression network analysis and applications to immunology. Bioinformatics. 2007;23(7):850–8.

107. Langfelder P, Horvath S. WGCNA: an R package for weighted correlation network analysis. BMC Bioinformatics. 2008;9(1).

108. Yip AM, Horvath S. Gene network interconnectedness and the generalized topological overlap measure. BMC Bioinformatics. 2007;8(1).

109. Harder AM, Willoughby JR, Ardren WR, Christie MR. Among-family variation in survival and gene expression uncovers adaptive genetic variation in a threatened fish. Mol Ecol. 2020;29(6):1035–49.

110. Huang DW, Sherman BT, Lempicki RA. Systematic and integrative analysis of large gene lists using DAVID bioinformatics resources. Nature Protocols. 2008;4(1):44–57.

111. Huang DW, Sherman BT, Lempicki RA. Bioinformatics enrichment tools: paths toward the comprehensive functional analysis of large gene lists. Nucleic Acids Research. 2009;37(1):1–13.

112. Raudvere U, Kolberg L, Kuzmin I, Arak T, Adler P, Peterson H, et al. g:Profiler: a web server for functional enrichment analysis and conversions of gene lists (2019 update). Nucleic Acids Research. 2019;47(W1):W191–W8.

113. Yao R, Jung D, Ge SX, Valencia A. ShinyGO: a graphical gene-set enrichment tool for animals and plants. Bioinformatics. 2020;36(8):2628–9.

114. Luo W, Pant G, Bhavnasi YK, Blanchard SG, Brouwer C. Pathview Web: user friendly pathway visualization and data integration. Nucleic Acids Research. 2017;45(W1):W501–W8.

